# Spatiotemporal Patterns of Gene Expression During Development of a Complex Colony Morphology

**DOI:** 10.1101/2024.09.14.613051

**Authors:** Gareth A. Cromie, Zhihao Tan, Michelle Hays, Amy Sirr, Aimée M. Dudley

## Abstract

Clonal communities of single celled organisms, such as bacterial or fungal colonies and biofilms, are spatially structured, with subdomains of cells experiencing differing environmental conditions. In the development of such communities, cell specialization is not only important to respond and adapt to the local environment but has the potential to increase the fitness of the clonal community through division of labor. Here, we examine colony development in a yeast strain (F13) that produces colonies with a highly structured “ruffled” phenotype in the colony periphery and an unstructured “smooth” phenotype in the colony center. We demonstrate that in the F13 genetic background deletions of transcription factors can either increase *(dig1*Δ, *sfl1*Δ) or decrease *(tec1*Δ) the degree of colony structure. To investigate the development of colony structure, we carried out gene expression analysis on F13 and the three deletion strains using RNA-seq. Samples were taken early in colony growth (day2), which precedes ruffled phenotype development in F13, and from the peripheral and central regions of colonies later in development (day5), at which time these regions are structured and unstructured (respectively) in F13. We identify genes responding additively and non-additively to the genotype and spatiotemporal factors and cluster these genes into a number of different expression patterns. We identify clusters whose expression correlates closely with the degree of colony structure in each sample and include genes with known roles in the development of colony structure. Individual deletion of 26 genes sampled from different clusters identified 5 with strong effects on colony morphology (*BUD8*, *CIS3*, *FLO11*, *MSB2* and *SFG1*), all of which eliminated or greatly reduced the structure of the F13 outer region.

## Introduction

Differentiation, the process by which cells specialize into distinct types, which can then be organized into larger-scale functional structures, is traditionally thought of as a phenomenon specific to multicellular organisms. In such organisms, differentiation increases fitness through structural and metabolic division of labor. Although the concept of differentiation is not meaningful in the context of isolated single-celled organisms, it is relevant to clonal communities of such organisms, such as bacterial or fungal colonies and biofilms. In these spatially-structured communities, cells in different regions may experience significantly different environments, including gas and nutrient availability, concentrations of waste and signaling products, exterior environmental challenges, and pH [1]. In these situations, cell specialization may not only benefit the individual that needs to respond and adapt to the local environment, but by structuring the community into cooperating domains of specialized cells, may improve the fitness of the community as a whole [2].

Some wild isolates of the budding yeast *Saccharomyces cerevisiae* produce highly structured “ruffled” colonies, rather than the unstructured “smooth” colony morphology typical of most laboratory strains. These structured colonies exhibit many of the features of biofilms, including production of an extracellular matrix, increased adherence, and localized expression of drug efflux pumps [3–5]. Often, these colonies are unstructured in the early stages of colony growth but become structured as the colony grows and ages. As part of this process, domains of specialized cells develop within ruffled colonies, with cells at the colony base forming pseudohyphae that invade the agar medium and anchor the colony, cells in peripheral regions upregulating drug efflux pumps and surface cells entering stationary phase, where they become more resistant to environmental insult [6]. Similarly, distinct gene expression profiles have been observed for the invasive cells within the agar, compared to the rest of the colony [7]. The molecular and regulatory pathways underlying ruffled colony formation overlap to a large extent with a group of related traits including mat formation and filamentous growth [8].

In a previous study [9], we identified six genes which repress ruffled colony morphology when overexpressed. We then deleted these genes in a strain background (F13) whose colonies on solid medium display a structured ruffled phenotype in the colony periphery and an unstructured smooth phenotype in the colony center. All six of the gene deletions promoted development of the ruffled morphology in the center of the F13 colonies, while maintaining the ruffled phenotype of the periphery. The six identified genes included the transcription factors *DIG1* and *SFL1*, that operate in the filamentation MAPK cascade and the PKA/cAMP pathways respectively and are known to have roles repressing ruffled colony formation and related phenotypes [10–12].

Here, we assess gene expression in F13 colonies, both before and after development of the structured outer region, and from both the outer ruffled and inner smooth regions of older colonies. We then use deletion of *SFL1* and *DIG1* to produce fully ruffled F13 colonies and demonstrate that deletion of the transcription factor *TEC1* results in completely smooth F13 colonies. The effect on gene expression of these genetic perturbations that increase and decrease the degree of colony morphology was then assessed. Our results identify spatiotemporal patterns of gene expression that reflect the development of a complex colony morphology.

## Materials and Methods

### Yeast Strains and Media

Unless noted, standard media and methods were used for growth and genetic manipulation of yeast [13] The strains of *S. cerevisiae* used in this study are listed in **Table 1**.

**Table 1.** Strains used in this study.

| Strain | Genotype | Origin |
| --- | --- | --- |
| YO780 (F13) | *MATα hoΔ0::hphMX4 SPS2:EGFP:NatMX* | [9] |
| YO1737 | *MATα hoΔ0::hphMX4 SPS2:EGFP:NatMX dig1Δ0::kanMX* | YO780 |
| YO1779 | *MATα hoΔ0::hphMX4 SPS2:EGFP:NatMX4 tec1Δ0::kanMX* | YO780 |
| YO1914 | *MATα hoΔ0::hphMX4 SPS2:EGFP:NatMX4 sfl1Δ0::kanMX* | YO780 |

### RNA Preparation and Sequencing

After 2 days of growth on YPD (2% glucose) plates at 30° C, whole colonies, arrayed in a “checkerboard” pattern [14], were harvested by scraping cells off the surface of the agar plate. To obtain sufficient quantities of RNA, 3-5 colonies were pooled for each sample, with three biological replicate samples taken of each strain/genotype. For the wild-type F13 strain, after 5 days of growth the smooth interior of the colony and the structured outside region were isolated separately, with inside and outside samples from 3-5 colonies pooled as above. For the deletion derivatives of F13, which form either fully smooth (*tec1*Δ) or fully ruffled (*sfl1*Δ, *dig1Δ*) colonies, inside and outside samples were taken to match the proportions taken from the F13 colonies.

Following extraction by hot acid phenol [15], total RNA from the pooled colonies was quantified by Bioanalyzer (Agilent). 5 µg of total RNA for each sample was then processed using the Tru-Seq stranded mRNA kit (Illumina) following manufacturer instructions. Individual sequencing libraries were pooled and analyzed by paired-end, 51-nucleotide read sequencing in one lane of an Illumina HiSeq 2000.

### Read-Pair Alignment

Read-pair alignment was carried out against the S288c reference (R64-1-1), with the FASTA and GFF files extended (**Files S1** and **S2**) to include ncRNAs and genes present in F45, a strain produced by the same cross that generated F13, but absent in S288c, as described previously [16]. Alignment was carried out using Bowtie2 (version 2.1.0) [17] with the parameters [-N 1 -I 50 -X 450 -p 6 --reorder -x -S] and allowing 1 mismatch per read.

For each strain, read alignments were then converted to gene counts using featureCounts (version 1.4.0) in the Subread package [18], with the parameters [-a -o -t gene –g ID –s 2 -T 1 -p -P -d 50 -D 450]. Reads were not filtered based on mapping quality, and thus we have been cautious in our interpretation of counts of genes that have paralogs with similar sequences, or which contain large regions of low sequence complexity. Read sequences are available from the Gene Expression Omnibus under accession GSE274952. Gene count tables are provided in **File S3**.

### Identification of Genes Responding to Genotype or Colony Age/Region

Analysis of gene expression was carried out using the edgeR [v. 3.6.8] [19] package for R [20] based on the tables of raw counts produced by featureCounts (**File S3**). Library sizes were first normalized using calcNormFactors (applying Trimmed Mean of M-values). Then the data was filtered to include only ORFs present in the S288c reference genome (genes with systematic names beginning with “Y”) and the counts for each gene in each library were converted to log_2_ counts per million reads, using a prior count of 20 to reduce the variance associated with low-expression genes. This produced a table with 4 genotypes, measured in triplicate, across 3 times/regions (day-2, day-5-inside and day-5-outside) giving a total of 36 normalized libraries (**File S3**). Two factor ANOVA models were then fit to this table with the first factor representing genotype (4 levels: *tec1*Δ, *wt*, *sfl1*Δ, *dig1*Δ) and the second factor representing the spatiotemporal variables (3 levels: day-2, day-5-inside and day-5-outside). Model fitting was performed twice, with the first ANOVA model allowing only an additive effect of the two factors (type II ANOVA) while the second ANOVA model included an interaction term between them (type III ANOVA). Multiple hypothesis correction was carried out for the p-value of the individual main and interaction terms in ANOVA separately using the Holm method [21]. We also tested each additive and interaction model against a null model (single mean) using ANOVA, and again multiple hypothesis corrected the results for each of the two models individually. For comparing the relative number of genes responding to the genotype and spatiotemporal factors, we used an adjusted p-value cutoff of 0.01 for each of the two terms in the additive ANOVA. To empirically confirm the stringency of our approach, we carried out one hundred permutations of the 36 sample measurements for each gene randomizing their sample label (i.e. the unique combinations of genotype spatiotemporal). We calculated the additive and interaction ANOVA p-values for each of these permutations and used these to calculate that our adjusted p-value cutoffs (p<0.01 versus null) for the additive and interaction ANOVAs correspond to empirical false discovery rates of 1.0 x 10^-05^ and 3.7 x 10^-05^, respectively.

We then identified genes with highly variable expression, having a minimum log_2_ expression variance of 0.2 across the whole dataset. Next, genes with an adjusted interaction term p-value of 0.01 or less from the interaction ANOVA were set aside as representing potential non-linear interactions between the two factors. From these, a final set of genes showing a strong non-additive response to both factors were then identified as those genes with a minimum r-squared of 0.9 in the interaction model. Among the remaining genes, those responding strongly and significantly to the individual factors were then identified from the first, additive ANOVA using a p-value cutoff of 0.01 (versus a null model) and a minimum r-squared cutoff of 0.7 for the full additive model, and a p-value cutoff of 0.01 for the factor of interest (all p-values were multiple hypothesis corrected, as described above).

### Characterization of Gene Expression Patterns

For examination of patterns of gene expression responding to genotype or to the spatiotemporal factor, or additively to both, the individual effect of each factor was isolated by normalizing for any effect of the other factor. This was done by setting the mean log expression of each gene within each level of the second factor to zero. After this normalization, the expression profiles of genes responding strongly and significantly to each factor underwent hierarchical clustering by gene. This was done using the pheatmap command of edgeR, with the expression of each gene centered and scaled to mean = 0 and variance = 1 and using the “complete linkage” clustering method.

For genes responding to the genotype factor, the three highest level clusters were extracted directly from the hierarchical clustering results. For genes responding to the spatiotemporal factor, eight major classes of gene expression were identified from visual inspection of the initial hierarchical clustering. These patterns were encoded as follows (2=high expression, 1=intermediate expression, 0=low expression) with each value applying to a single level of the spatiotemporal factor (day-2, day-5-inside, day-5-outside; each containing three replicates of the four genotypes): C1=(2,0,0), C2=(0,2,2), C3=(2,2,0), C4=(0,0,2), C5=(2,1,0), C6=(0,1,2), C7=(0,2,0), C8=(2,0,2). Each gene was then assigned to the class whose template pattern showed the highest correlation with the expression pattern of the gene. Expression heatmaps for all individual classes were produced using the pheatmap command, from the pheatmap package [22] in R.

### Functional Enrichment of Gene Lists

Functional enrichment of *S. cerevisiae* gene lists was performed using g:Profiler [23] with a multiple hypothesis corrected (g:SCS) threshold of p<0.01.

### Availability of Data and Material

The datasets generated during and/or analyzed in the current study are available in the Gene Expression Omnibus (GEO) under accession GSE274952 The script used to carry out the data processing and analysis is given in **File S4**.

## Results

### Strain F13 is a Model for Detecting both Increases and Decreases in the Degree of Colony Structure

The haploid budding yeast strain F13, derived from a cross between a sake-brewing strain and an Ethiopian white tecc strain [24], displays an unusual pattern of colony development on solid medium. Initially, the colony is unstructured, similar to common laboratory strains, but after three days of growth develops a clearly-defined, highly-structured outer ring, with a smooth, unstructured center (**Figure 1**).

**Figure 1.**
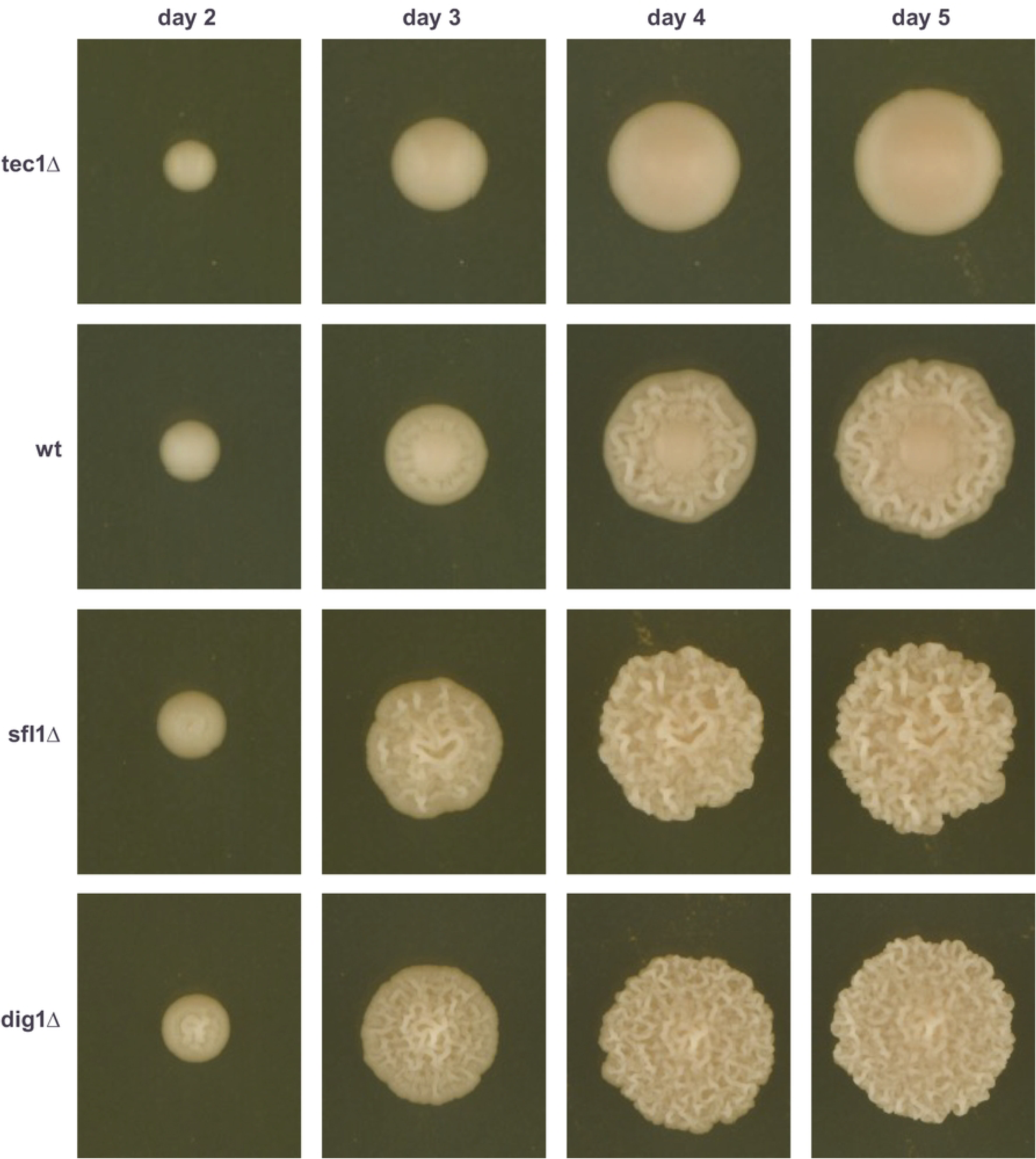
Morphology of F13 colonies as they develop over time and effect on colony morphology of deleting *TEC1*, *SFL1* and *DIG1*.

The unusual colony morphology of late-stage F13 colonies, having both smooth and structured zones, suggested that F13 might allow detection both of perturbations increasing, and perturbations decreasing, the degree of complex colony morphology. We previously demonstrated that genetic perturbations expected to increase colony structure cause F13 to form fully structured colonies, with no smooth center at late time points [9] (**Figure 1**). Specifically, we deleted genes encoding two known inhibitors of complex colony morphologies, Dig1 and Sfl1, that operate in the filamentation MAPK cascade and the PKA/cAMP pathways respectively [10–12]. Colonies of these genotypes have already begun to develop colony structure by day-2, when F13 colonies are still fully smooth (**Figure 1**). Here, we additionally tested the ability of F13 to respond to genetic perturbations that decrease colony structure, by deleting *TEC1*, a gene that also encodes a member of the filamentation MAPK cascade, but which promotes complex colony morphology [12]. Deletion of *TEC1* produced colonies that remained fully smooth throughout their development (**Figure 1**). Therefore, it appears that strain F13 is indeed a model allowing detection both of perturbations increasing, and perturbations decreasing, colony structure.

### Patterns of Gene Expression in F13

To explore the patterns of gene expression associated with the regional development of colony morphology over time in strain F13 we carried out RNA-seq on complete F13 colonies from day-2 of growth (when colonies of all genotypes are still unstructured) and on the outer (structured) and inner (unstructured) regions of 5-day-old F13 colonies. RNA-seq was also carried out on day-2 and day-5 colonies of strains with deletions of *DIG1*, *SFL1* or *TEC1*. Although colonies with these deletion genotypes were either fully smooth or fully structured at day 5, inner and outer portions of those colonies, chosen to match the relative proportions observed in wild type F13, were isolated and analyzed separately (**Materials and Methods)**. Therefore, day-2, day-5 (outside) and day-5 (inside) samples were isolated for all four genotypes. Three biological replicates were analyzed for each of these sample types.

To identify the major patterns of gene expression relating to genotype, colony age and colony region, we carried out a two-way additive ANOVA on the gene expression data (normalized log_2_ counts per million reads) (**Materials and Methods**). The first factor represented the genotype and had 4 levels: *tec1*Δ, wild-type F13 (WT), *sfl1*Δ and *dig1*Δ. The second factor captured the spatiotemporal variables and had 3 levels: day-2 (whole colony), day-5 (outside) and day-5 (inside). We identified 2,978 genes that showed a significant response to the two factors under an additive model (type II Anova p<0.01, after multiple hypothesis correction). Using permutation testing (**Materials and Methods**), we determined that our chosen significance threshold corresponds to an empirical false discovery rate of 1.0 x 10^-5^. Among the set of genes, examination of the p-values associated with the two factors indicated that many more genes (2,772 vs 344) showed a significant (p<0.01, after multiple hypothesis correction) change in expression in response to the spatiotemporal factor than to the genotype factor (**Figure 2**). A total of 221 genes showed a significant additive response to both factors (see below).

**Figure 2.**
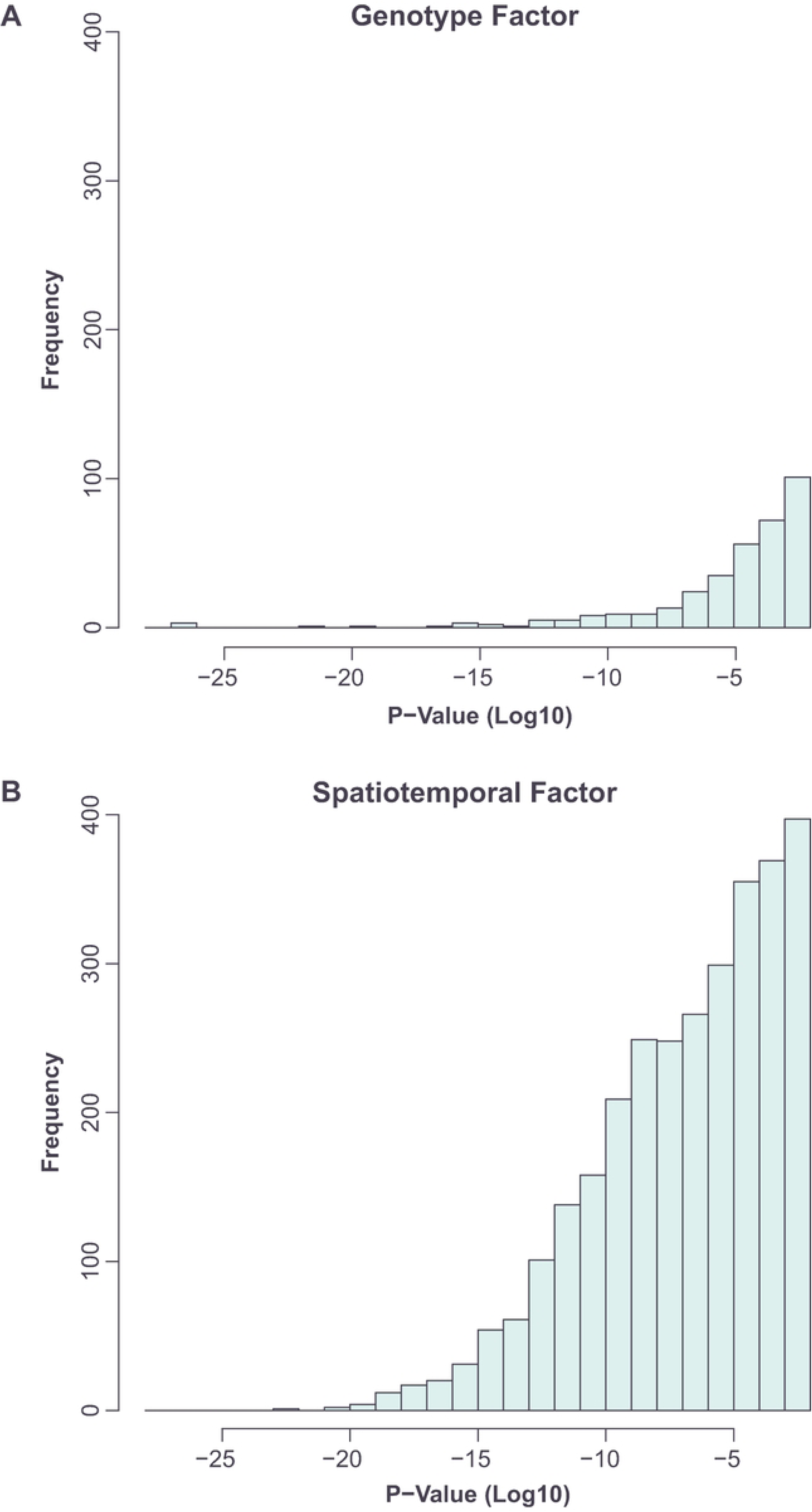
In an additive model, a smaller number of genes show a significant (p<0.01, multiple hypothesis corrected) response to the genotype factor (A) than to the spatiotemporal factor (B).

To isolate the major genotype-dependent and spatiotemporal patterns of gene expression we normalized our expression data for the effect of one factor and carried out hierarchical clustering to identify major patterns of gene expression associated with the remaining factor in isolation (for genes significantly responding to that factor, **Materials and Methods**). We focused on genes with the strongest and clearest additive response to the two factors, i.e. genes having a high degree of variation in expression within the experiment (variance of normalized log counts > 0.2) that was well explained by the additive model (r^2^>0.7). A small number of genes whose expression was better explained by a non-additive interaction between the two factors (**Materials and Methods**) were excluded at this point for later analysis (see below).

### Genotype-Dependent Patterns of Gene Expression

After normalizing for the spatiotemporal factor and carrying out hierarchical clustering (**Materials and Methods**), three major patterns of gene expression were visible, among genes with statistically significant responses to the genotype factor (**Figure 3**).

**Figure 3.**
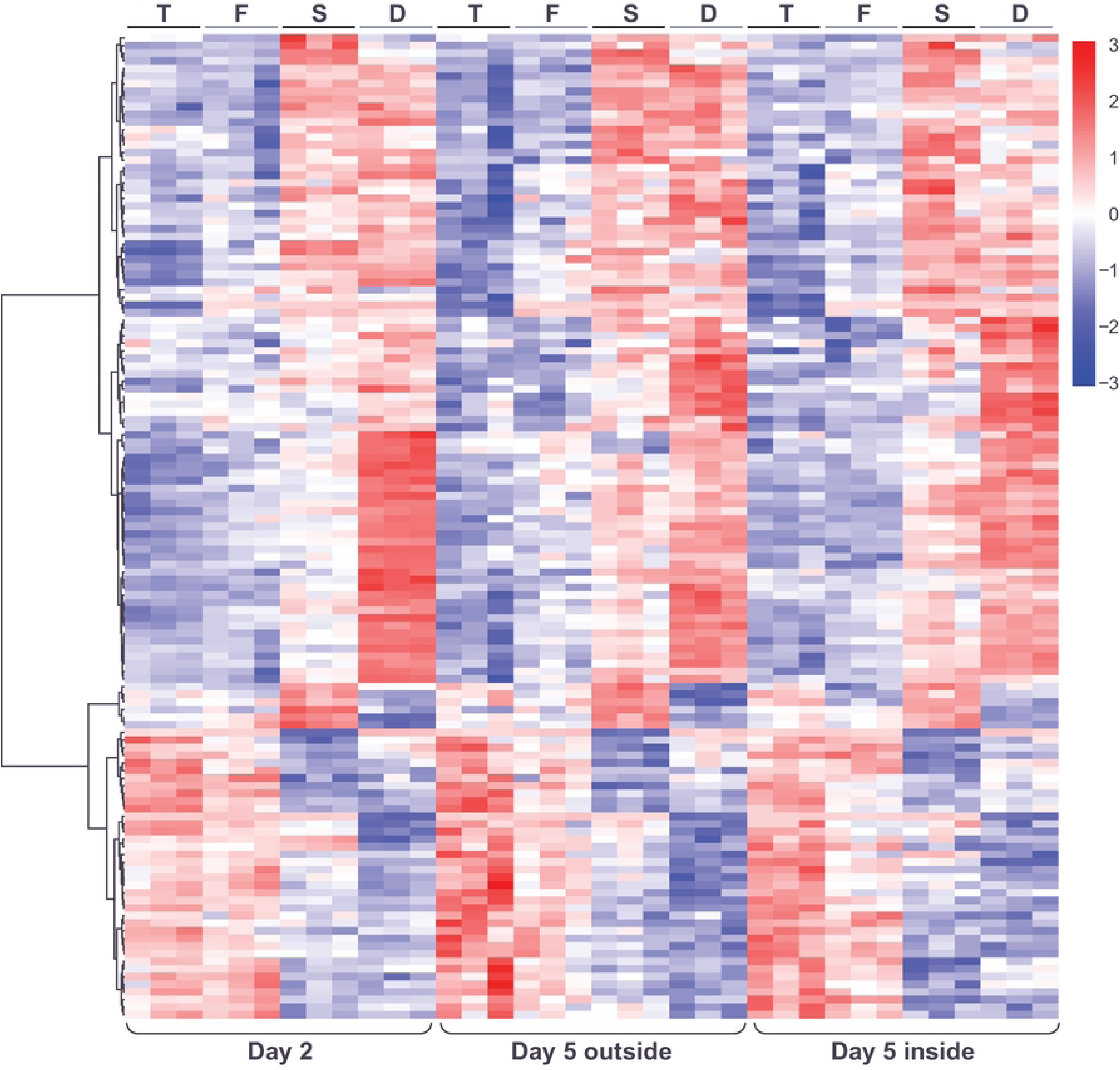
Heatmap showing hierarchical clustering of genes with significant changes in expression in response to genotype (after normalization for the spatiotemporal factor), across the three spatiotemporal conditions and three replicates. Color in heatmaps reflects log_2_ normalized expression for each gene (red high, blue low). Heatmap genotypes are indicated as T=*tec1*Δ, F=WT F13, S=*sfl1*Δ, D=*dig1*Δ.

The first pair of clusters (G1 and G2) identified genes with mirror patterns of gene expression in the order *tec1*Δ, WT*, sfl1*Δ, *dig1*Δ, with *dig1*Δ showing the highest expression in cluster G1 and the lowest expression in cluster G2 (**Figure 4**). The order of these genotypes corresponds to the associated degree of colony morphology seen at day 5, with *tec1*Δ fully smooth, F13 WT intermediate and *sfl1*Δ and *dig1*Δ fully ruffled, i.e. the expression level of these genes correlates (positively in cluster G1 and negatively in cluster G2) with the degree of colony structure associated with each genotype. Genes in cluster G1 are significantly enriched (p<0.01) for a number of GO terms, including “fungal-type cell wall” (GO:0009277; p=2.71e-04), “extracellular region” (GO:0005576; p=7.78e-04) and “regulation of establishment or maintenance of cell polarity” (GO:0032878; p=5.67e-04), and for targets of the transcription factor Tec1 (TF:M01810_0; p=3.38e-07) (**Table S1**). Genes in cluster G2 are significantly enriched for several GO terms including “extracellular region” (GO:0005576; p=1.78e-11), “fungal-type cell wall” (GO:0009277; p=1.72e-09), “cell wall organization” (GO:0071555; p=1.42e-05), “hydrolase activity, hydrolyzing O-glycosyl compounds” (GO:0004553; p=7.59e-05) and “side of membrane” (GO:0098552; p= 2.51e-04) and for targets of the transcription factor Matalpha2-Mcm1 (TF:M08191; p=1.58e-04) (**Table S2**).

**Figure 4.**
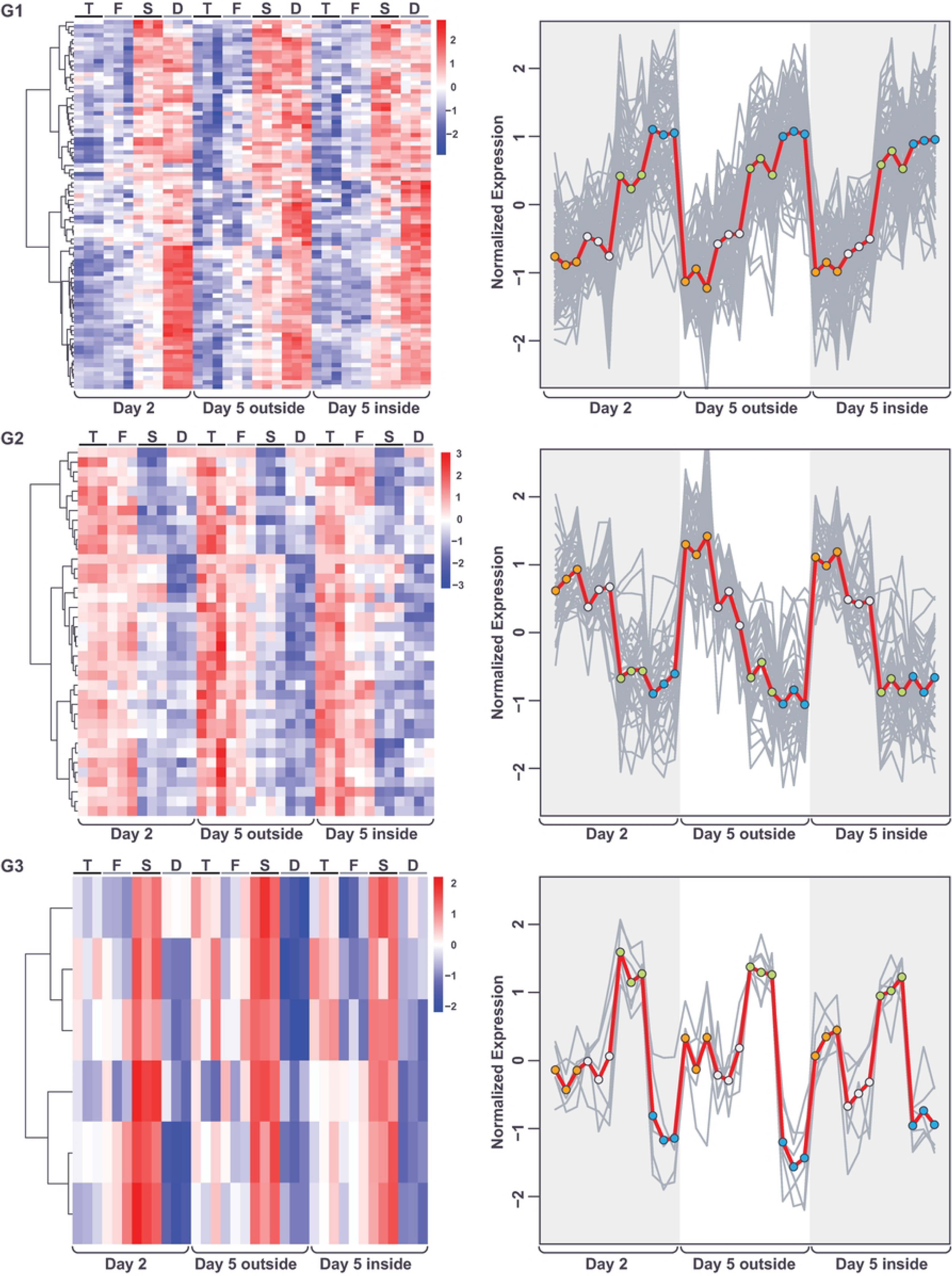
Gene expression patterns (after normalization for spatiotemporal factor) within each of the 3 genotype clusters displayed as heatmaps (left) and expression traces (right) of all genes within each cluster (grey lines). In the expression traces, the mean expression of the genes in the cluster is highlighted in red line with circles reflecting genotype (orange=*tec1Δ*, grey=*wt*, green=*sfl1Δ*, blue=*dig1Δ*). Color in heatmaps reflects log_2_ normalized expression for each gene (red high, blue low) and genotypes are indicated as T=*tec1*Δ, F=WT F13, S=*sfl1*Δ, D=*dig1*Δ.

The final cluster (G3) identified a group of only 6 genes with highest expression in *sfl1*Δ and lowest expression in *dig1*Δ. These genes are significantly enriched for several GO terms, including “water channel activity” (GO:0015250; 5.62e-04) and water transport (GO:0006833; 2.63e-03) (**Table S3**).

### Spatiotemporal Patterns of Gene Expression

After normalizing for genotype and carrying out hierarchical clustering (**Materials and Methods**), 4 major patterns of gene expression were visible, among genes with statistically significant responses to the spatiotemporal factor. These four major patterns could be grouped into two sets of “mirror” pairs. The first pair consisted of one group of genes with high expression in the day-5-outside sample and low expression in the other two samples, and one group of genes with the reverse pattern, having lowest expression in the day-5-outside sample (**Figure 5**). Both of these groups were small. The second pair consisted of one large group of genes with high expression at day-2 and low expression in the day-5-inside sample, and another large group showing the reverse pattern (day-2 low, day-5-inside high) (**Figure 5**). Within this second pair, a continuum of expression levels was observed in the day-5-outside sample, therefore we further split these two sets of genes based on whether the day-5-outside expression level was intermediate between the day-2 and day-5-inside samples, or whether it more closely matched either the day-2 or day-5-inside expression level (**Materials and Methods**). This gave us the 8 classes shown in **Figures 6 and 7**, consisting of four pairs of “mirror” patterns.

**Figure 5.**
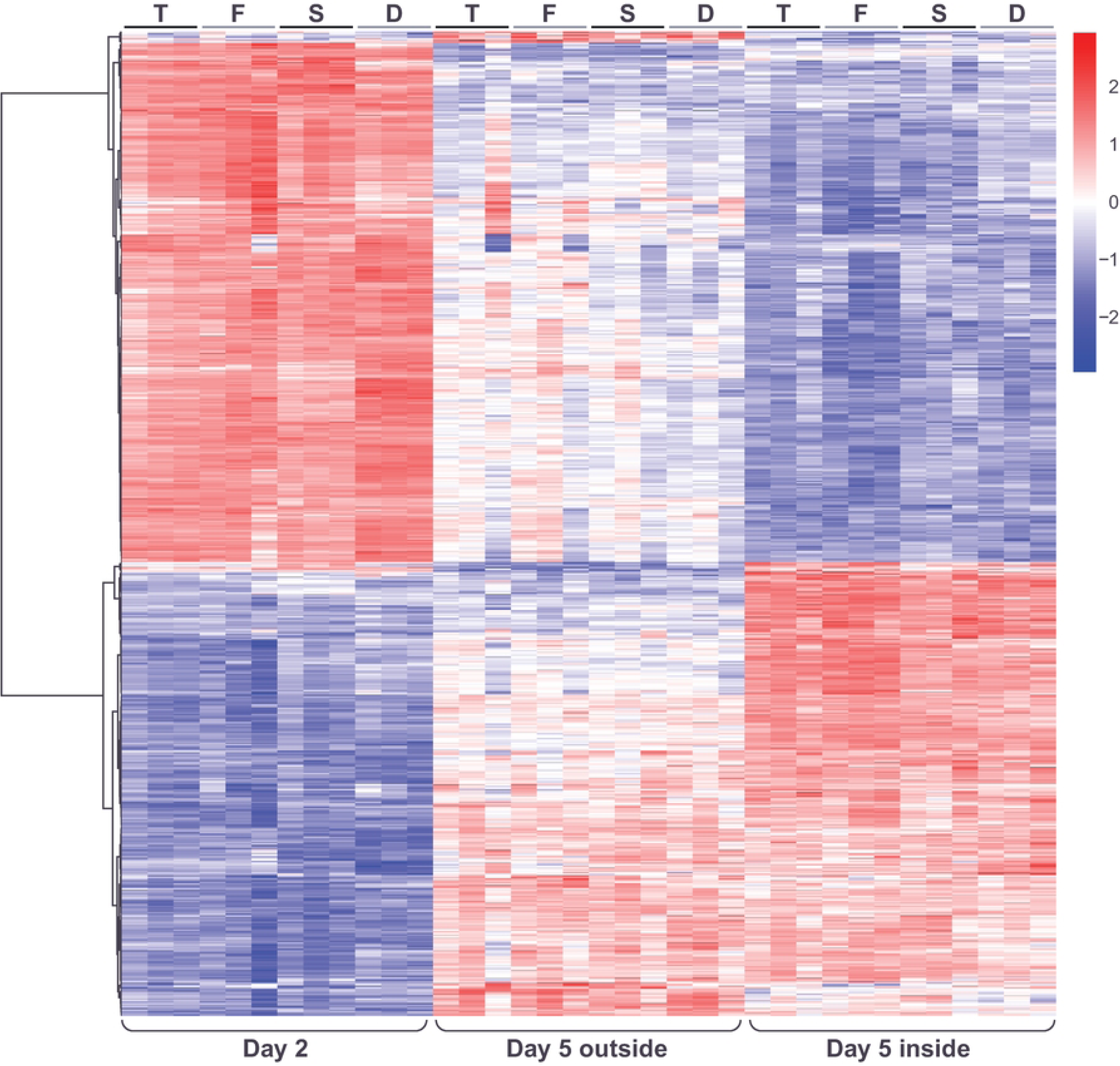
Heatmap showing hierarchical clustering of genes with significant changes in expression in response to the spatiotemporal factor (after normalization for genotype), across the four genotypes and three replicates. Color in heatmaps reflects log_2_ normalized expression for each gene (red high, blue low) and genotypes are indicated as T=*tec1*Δ, F=WT F13, S=*sfl1*Δ, D=*dig1*Δ.

**Figure 6.**
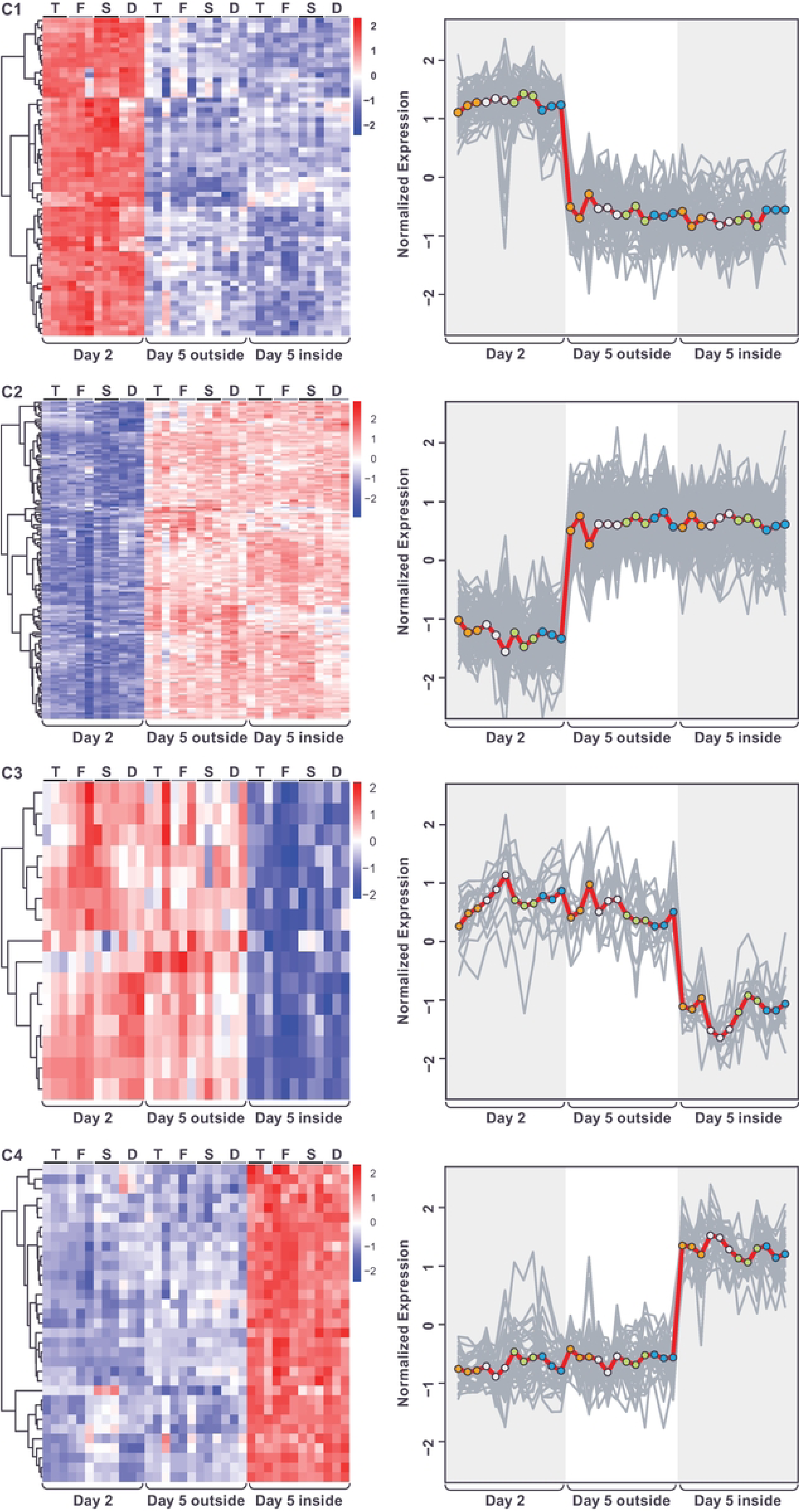
Gene expression patterns (after normalization for genotype) within each of the first four spatiotemporal clusters displayed as heatmaps (left) and expression traces (right) of all genes within each cluster (grey lines). In the expression traces, the mean expression of the genes in the cluster is highlighted in red line with circles reflecting genotype (orange=*tec1*Δ, grey=WT F13, green=*sfl1*Δ, blue=*dig1*Δ). Color in heatmaps reflects log_2_ normalized expression for each gene (red high, blue low) and genotypes are indicated as T=*tec1*Δ, F=WT F13, S=*sfl1*Δ, D=*dig1*Δ.

**Figure 7.**
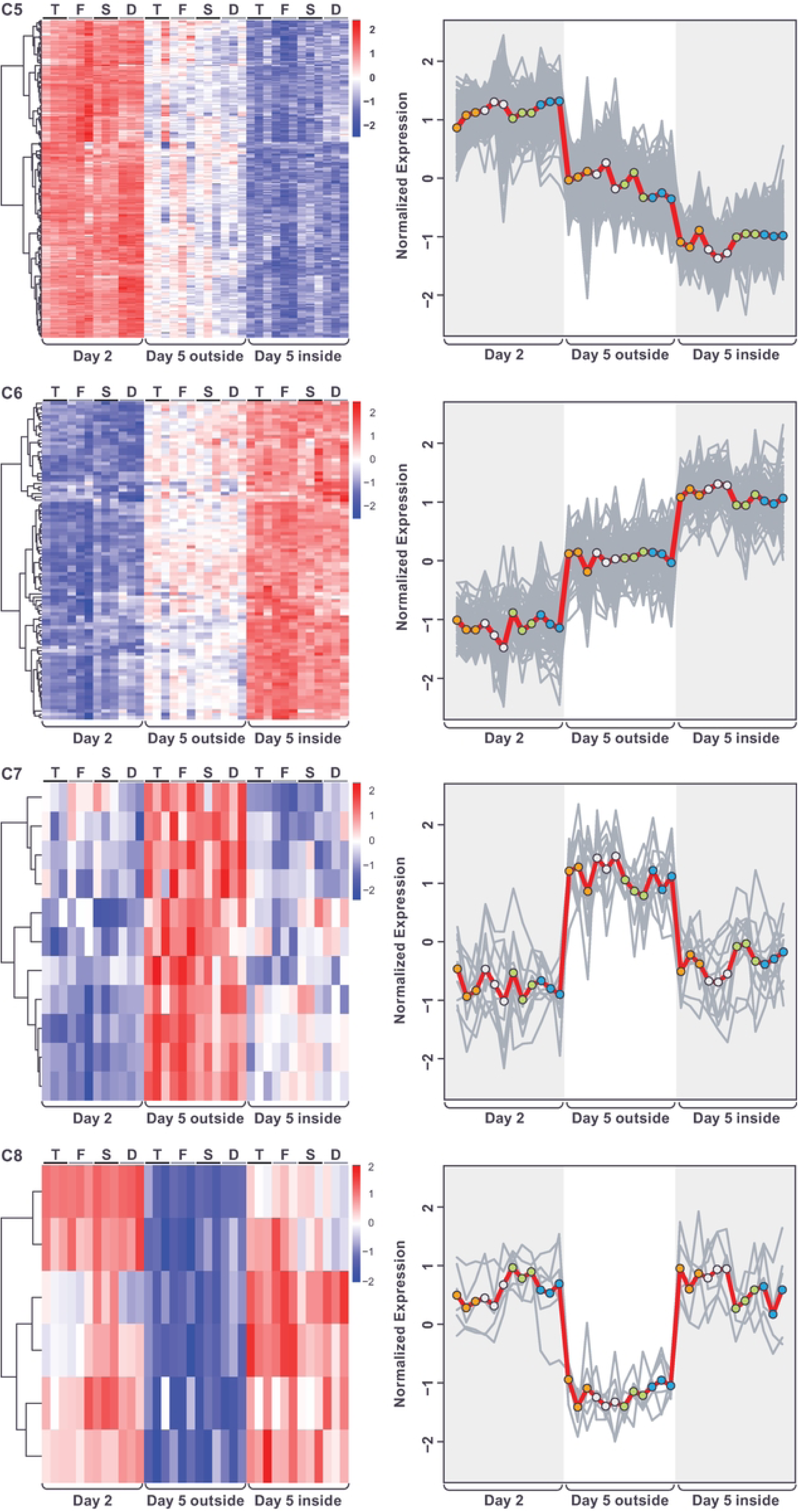
Gene expression patterns (after normalization for genotype) within each of the second four spatiotemporal clusters displayed as heatmaps (left) and expression traces (right) of all genes within each cluster (grey lines). In the expression traces, the mean expression of the genes in the cluster is highlighted in red line with circles reflecting genotype (orange=*tec1*Δ, grey=WT F13 green=*sfl1*Δ, blue=*dig1*Δ). Color in heatmaps reflects log_2_ normalized expression for each gene (red high, blue low) and genotypes are indicated as T=*tec1*Δ, F=WT F13, S=*sfl1*Δ, D=*dig1*Δ.

The first pair of these “mirror” clusters (C1 and C2) identified genes with differential expression between day-2 and day-5, with little difference in expression within the day-5 colony between the outside and inside regions, i.e. these genes seem to be responding only to colony age/size. In cluster C1, genes were repressed at day-5 while in cluster C2 they were induced (**Figure 6**). Genes in cluster C1 are significantly enriched for a number of GO terms that include “extracellular region” (GO:0005576; p=4.23e-06), “fungal-type cell wall” (GO:0009277; p= 1.25e-05) and “side of membrane” (GO:0098552; p=7.91e-05 (**Table S4**), while genes in cluster C2 are significantly enriched for several GO terms, including “cellular component assembly involved in morphogenesis” (GO:0010927; p=6.06e-05), “fungal-type cell wall” (GO:0009277; p=1.86e-03), “prospore membrane” (GO:0005628; p=6.33-03) and “glyoxalase III activity” (GO:0019172; p=9.42e-03) (**Table S5**).

The second pair of clusters (C3 and C4), identified genes with similar expression levels between the day-2 and the day-5-outside samples but with a different level of expression in the day-5-inside sample, i.e. reflecting expression changes specific to the inside of the day-5 colonies. In cluster C3 the genes were repressed in the day-5-inside sample and in cluster C4 they were induced (**Figure 6**). No significant GO or KEGG term enrichment was seen for genes in cluster C3 (**Table S6**). Genes in cluster C4 were significantly enriched for a number of GO terms including “monocarboxylic acid metabolic process” (GO:0032787; p=2.61e-07), “cellular lipid metabolic process” (GO:0044255; p=2.59e-06) “C-acyltransferase activity” (GO:0016408; p=1.00e-03), “ammonium transmembrane transporter activity” (GO:0008519; p=1.00e-03) and “fatty acid synthase complex” (GO:0005835; p=3.48e-03) (**Table S7**). The third pair of clusters (C5 and C6) identified genes where the level of expression in the day-5-outside sample was intermediate between that of the day-2 and day-5-inside samples. In cluster C5, expression was lowest in the day-5-inside sample while in cluster C6 expression was highest in that sample (**Figure 7**). Genes in cluster C5 were enriched for GO terms that include “cytoskeleton” (GO:0005856; p=2.09e-12), “amino acid biosynthetic process” (GO:0008652; p=3.42e-12), “cellular bud” (GO:0005933; p=2.96e-11) and “heterocyclic compound binding” (GO:1901363; p=2.80e-06) and the KEGG term “Biosynthesis of amino acids” (KEGG:01230; p=4.54e-15) (**Table S8**). Genes in cluster C6 were enriched for GO terms including “sorbitol transport” (GO:0015795; p= 2.37e-05), “monocarboxylic acid metabolic process” (GO:0032787; p=7.41e-05) and “peroxisome” (GO:0005777; p=3.28e-04) (**Table S9**).

The final pair of clusters (C7 and C8) identified the small number of genes where the level of expression in day-2 and day-5-inside samples was similar, but this differed from expression in the day-5-outside sample, i.e. identifying expression specific to the outside region of the day-5 colonies. In cluster C7 (**Table S10**), expression was high in the day-5-outside sample while in cluster C8 (**Table S11**) expression was low in that sample (**Figure 7**). No significant GO or KEGG term enrichment was seen for genes in either cluster.

### Genes Responding Additively to both Genotype and Spatiotemporal Factors

We next examined those genes whose expression responded significantly (and additively) to both the genotype and spatiotemporal factors. The number of genes responding to both factors was much larger than expected by chance, with 616 genes responding to the spatiotemporal factor, 129 genes responding to genotype, and 103 genes responding to both (Fisher’s Exact Test: one tailed p=4.73e-84; 6575 genes total). In addition, the distribution of counts (**Table 2**) across the clusters of the two factors was also non-independent (Fisher’s Exact Test: two tailed p=2.05e-08).

**Table 2.** Overlap between genes in spatiotemporal clusters (C1-8) and genotype clusters (G1-G3).

| Cluster | G1 | G2 | G3 |
| --- | --- | --- | --- |
| C1 | 0 | 11 | 0 |
| C2 | 39 | 4 | 2 |
| C3 | 5 | 1 | 0 |
| C4 | 1 | 0 | 0 |
| C5 | 11 | 12 | 0 |
| C6 | 8 | 2 | 1 |
| C7 | 4 | 0 | 1 |
| C8 | 0 | 1 | 0 |

For genes in cluster G1 (gene expression correlates positively with degree of fluffiness of genotype) whose expression also responded to the spatiotemporal factor, an over-representation was seen in cluster C2 (low expression in day-2 and high expression in both day-5 samples) and an under-representation was seen in cluster C1 (high expression in day-2 and low expression in both day-5 samples), compared to expectation under independence. In fact, out of all 85 genes in cluster G1 (regardless of any response to the spatiotemporal factor) a total of 39 (46%) were also placed in cluster C2. Taken together, these data suggest that there is a significant overlap between genes whose expression correlates positively with more structured genotypes and genes whose expression is higher in day-5, when ruffled morphology is most strongly developed, than in day-2, when it is weak or has not yet developed.

The exact opposite pattern was seen for genes in cluster G2 (gene expression correlates negatively with degree of fluffiness of genotype) whose expression also responded to the spatiotemporal factor. For these genes, an under-representation was seen in cluster C2 (low expression in day-2 and high expression in both day-5 samples) and an over-representation was seen in cluster C1 (high expression in day-2 and low expression in both day-5 samples), compared to the expectation under independence. That is, there is a strong overlap between genes whose expression correlates negatively with more structured genotypes and genes whose expression is higher in day-2 than in day-5.

Taken together, the overlap patterns seen with clusters G1 and G2 suggest the existence of a reciprocal relationship between expression in the structured vs smooth genotypes and expression in day-5 versus day-2. These patterns suggest that these two sets of genes (G1+C2 and G2+C1) might include genes important for the development of ruffled versus smooth colony morphologies.

The 39 genes assigned to both clusters G1 and C2 did not show any significant GO term enrichment (**Table S12**) while the G2+C1 group (**Table S13**) was significantly enriched for several related GO terms, including “extracellular region” (GO:0005576; p=1.04e-04) as well as “fungal-type cell wall” (GO:0009277; p= 1.75e-04), and “side of membrane” (GO:0098552; p=1.65e-03).

### Genes Responding Non-Additively to both Genotype and Spatiotemporal Factors

Having completed our analysis of genes responding additively to the genotype and spatiotemporal factors, we examined those genes whose expression showed a significant non additive response to the two factors (**Materials and Methods**). These genes had been excluded from the preceding analyses. As a group (**Table S14**), these genes show strong enrichment for the GO terms “cell periphery” (GO:0071944; p=1.64e-10) and “extracellular region” (GO:0005576; p=1.14e-08). Hierarchical clustering of these genes identified several distinct expression patterns, which we extracted as 9 clusters (**Figure 8** and **Table S11**).

**Figure 8.**
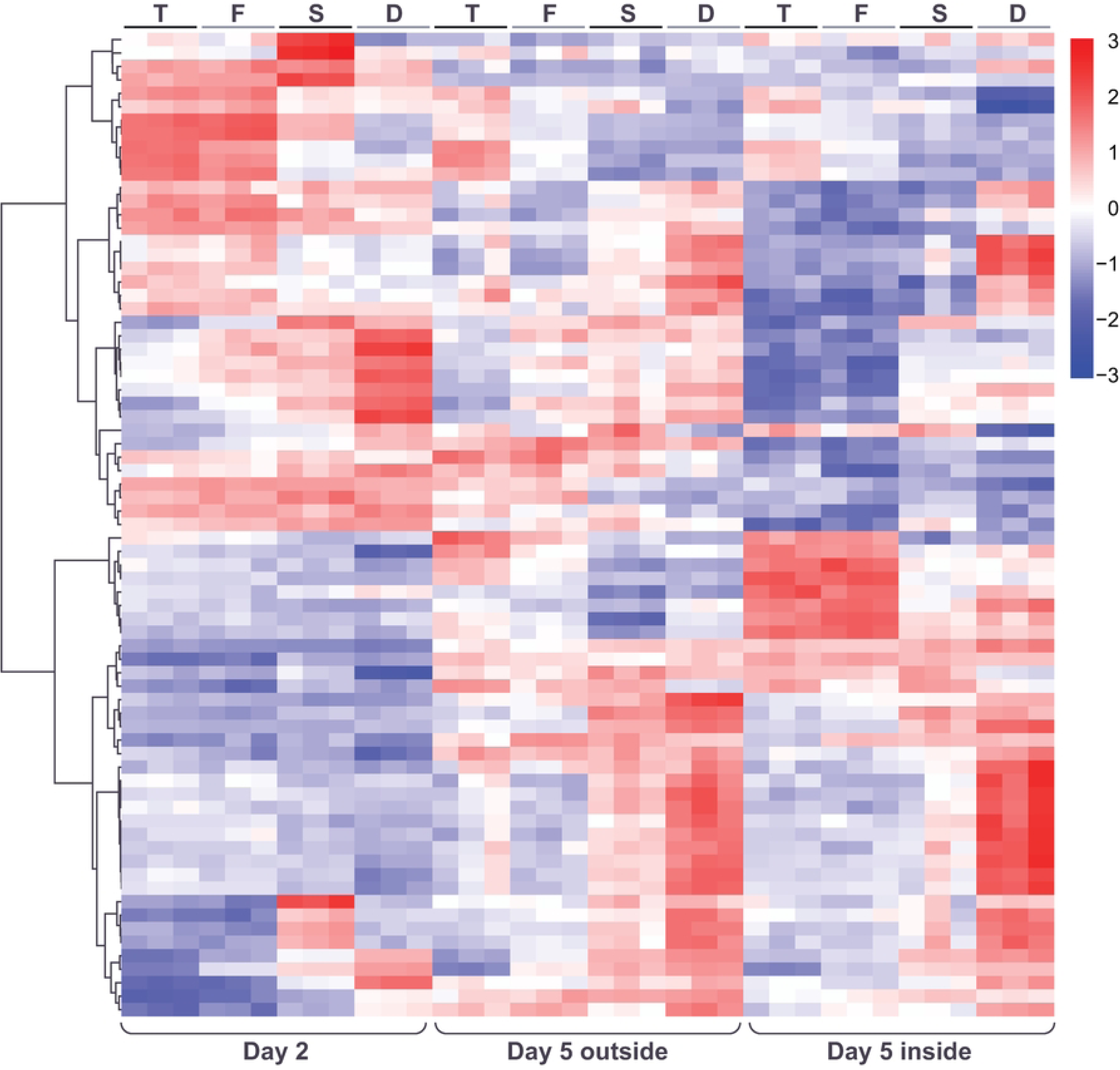
Heatmap showing hierarchical clustering of genes whose expression indicates a significant interaction between the genotype and spatiotemporal factors. Color in heatmaps reflects log_2_ normalized expression for each gene (red high, blue low) and genotypes are indicated as T=*tec1*Δ, F=WT F13, S=*sfl1*Δ, D=*dig1*Δ.

One of these clusters (cluster I9), with eight members, included several genes that are known to have important roles in the development of complex colony morphologies. These genes are *FLO11*, encoding a cell wall flocculin believed to play a critical mechanistic role in pseudohyphal and filamentous growth and in the development of structured colonies [12, 25, 26], *PHD1*, encoding a transcriptional enhancer that promotes pseudohyphal growth (via mechanisms such as regulating *FLO11* expression level) [27, 28], and *MSB2*, encoding a cell wall mucin protein that promotes filamentous growth via the “filamentous” MAPK pathway [29]. These three genes either encode cell wall components or regulate cell wall processes and several other members of this cluster also have cell wall roles, with the paralogs *SVS1* and *SRL1* encoding cell wall proteins [30] and *WSC2* encoding a signal transducer in the stress-activated *PKC1*-*MPK1* signaling pathway which helps maintain cell wall integrity [31]. The final two genes are *SIM1*, encoding a SUN family protein that may participate in DNA replication [32], and *TOS7*, encoding a protein with likely roles in secretion and cell wall organization [33]. The genes in cluster I9 are enriched for several GO terms including “cell surface” (GO:0009986; p=7.51e-05), “cell periphery” (GO:0071944; p=4.35e-04) and “site of polarized growth” (GO:0030427; p=5.12e-04) and the KEGG pathway “MAPK signaling pathway – yeast” (KEG:04011; p=1.72e-03) (**Table S15**).

The pattern of gene expression in cluster I9 in the day-5 samples correlates closely and positively with the degree of ruffled colony morphology in those samples (**Figure 9**). Expression levels are low in the *tec1*Δ day-5-outside sample and the *tec1*Δ and WT day-5-inside samples, which are all smooth. Conversely, expression is higher in the WT, *sfl1*Δ and *dig1*Δ day-5-outside samples, and the *sfl1*Δ and *dig1*Δ day-5-inside samples, all of which are ruffled. Notably, gene expression in the day-5 outside samples differs between *tec1*Δ (smooth) and F13 WT (ruffled) while expression is similar between these genotypes in the day-5 inside samples (both smooth). Expression in the day-2 samples correlates positively with the degree of fluffiness associated with each genotype at day-5, increasing in the order *tec1*Δ, F13 WT, *sfl1*Δ and *dig1*Δ, similar to cluster G1 in the analysis of the additive ANOVA.

**Figure 9.**
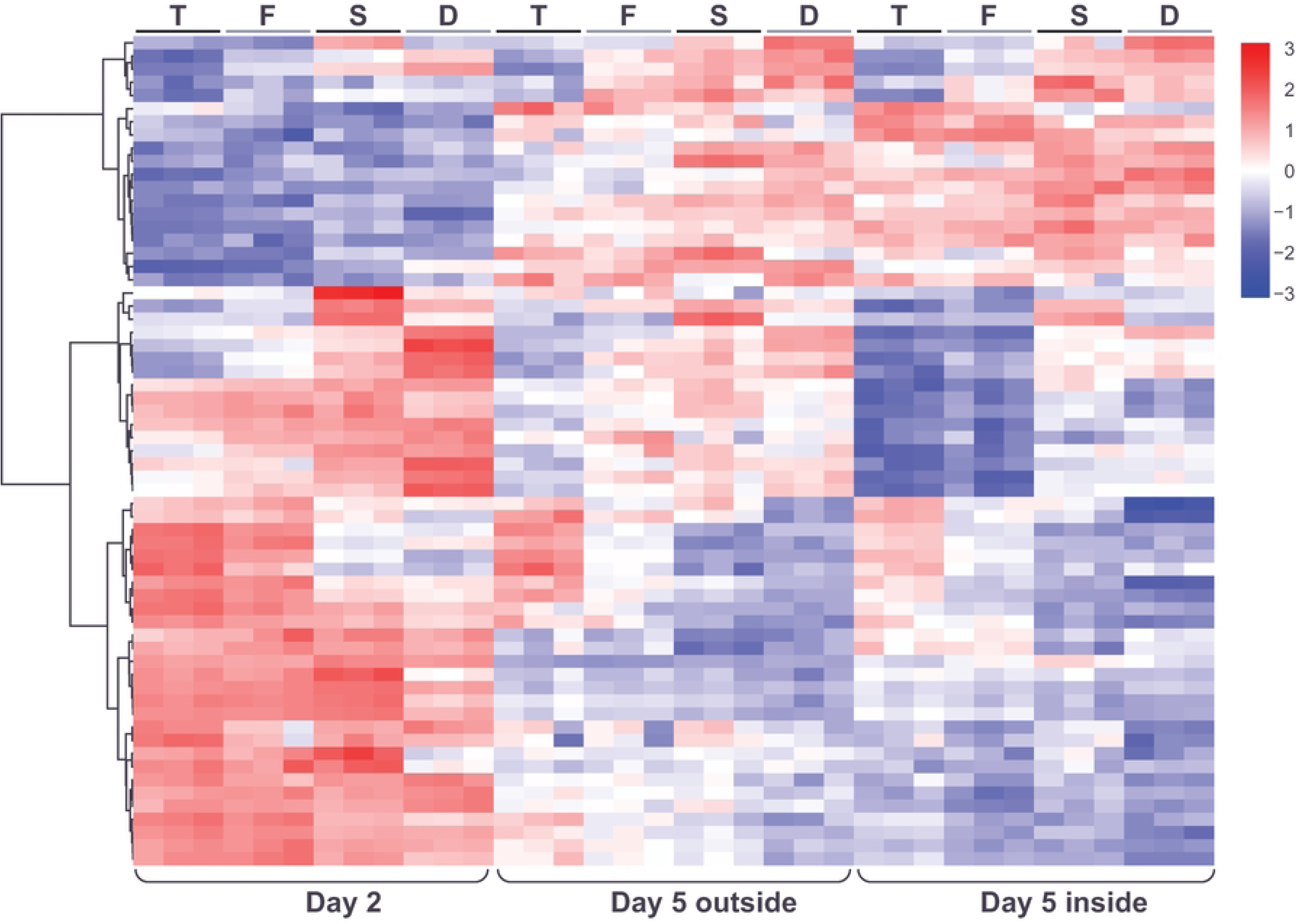
Gene expression patterns within the spatiotemporal/genotype interaction clusters I1 and I9 displayed as heatmaps (left) and expression traces (right) of all genes within each cluster (grey lines). In the expression traces, the mean expression of the genes in the cluster is highlighted in red line with circles reflecting genotype (orange=*tec1*Δ, grey=WT F13, green=*sfl1*Δ, blue=*dig1*Δ). Color in heatmaps reflects log_2_ normalized expression for each gene (red high, blue low) and genotypes are indicated as T=*tec1*Δ, F=WT F13, S=*sfl1*Δ, D=*dig1*Δ.

Cluster I1 shows a pattern of gene expression that is essentially the inverse of cluster I9 across the day-5 genotypes and colony regions (**Figure 9**). However, expression in the day-2 samples varies little across the genotypes. Cluster I1 (**Table S16**) consists of 8 genes (*GDH3*, *MUP3*, *SYG1*, *SNZ1*, *FDH1*, *SSU1*, *OPT2* and *YHR137C-A)*, showing no enrichment for any GO or KEGG pathways.

### Changes in the Expression of Genes Encoding Extracellular Proteins in Response to the Genotype and Spatiotemporal Factors

Several of the gene expression clusters that we identified in our analysis showed enrichment for genes encoding extracellular and cell wall proteins. This included the set of all genes showing a non-additive interaction between the genotype and spatiotemporal factors. In addition, several of the paired clusters that show inverse patterns of gene expression (e.g. C1 versus C2 and G1 versus G2) showed enrichment for extracellular genes in both of the clusters, despite their opposite patterns of expression. To look into the behavior of extracellular genes more directly we carried out cluster analysis of all genes with the GO terms (GO:0005576 “extracellular region”, GO:0009277 “fungal-type cell wall”, GO:0009986 “cell surface”, GO:0031505 “fungal-type cell wall organization”) and whose expression responded significantly to the additive model using the genotype and spatiotemporal factors described earlier or showed a significant non-additive response to the two factors (**Materials and Methods**).

Based on this cluster analysis, it appears that the extracellular genes can be classified as responding mostly to the spatiotemporal factor, or mostly to the genotype factor (**Figure 10** and **Table S17**). Among the genes responding mainly to the spatiotemporal factor, two “reflected” patterns were observed with genes either showing highest expression at day-2 and lowest expression in the day-5-inside sample (Cluster E1), or vice versa (Cluster E2) (**Figure 11**). Genes encoding glycosylated cell surface proteins were observed in both of these clusters (E1: *DAN1*, *FLO5*, *FLO9*, TIR1, *TIR2*, *TIR3*, *TIR4*; E2: *CWP2*, *CWP1*, *FIT2*, *PAU24*, *PHO5*, *PIR3*, *SED1*) (**Table S17**). In contrast, cluster E2 contained several spore wall genes (*GAS4*, *SPO11*, *SPS22*) and the two genes encoding a-factor (*MFA1*, *MFA2*) while cluster E1 contained genes encoding several enzymes involved in cell wall modification including glucanases (*EGT2*, *EXG2*, *SUN4*) and a chitin transglycosylase (*UTR2*) (**Table S17**).

**Figure 10.**
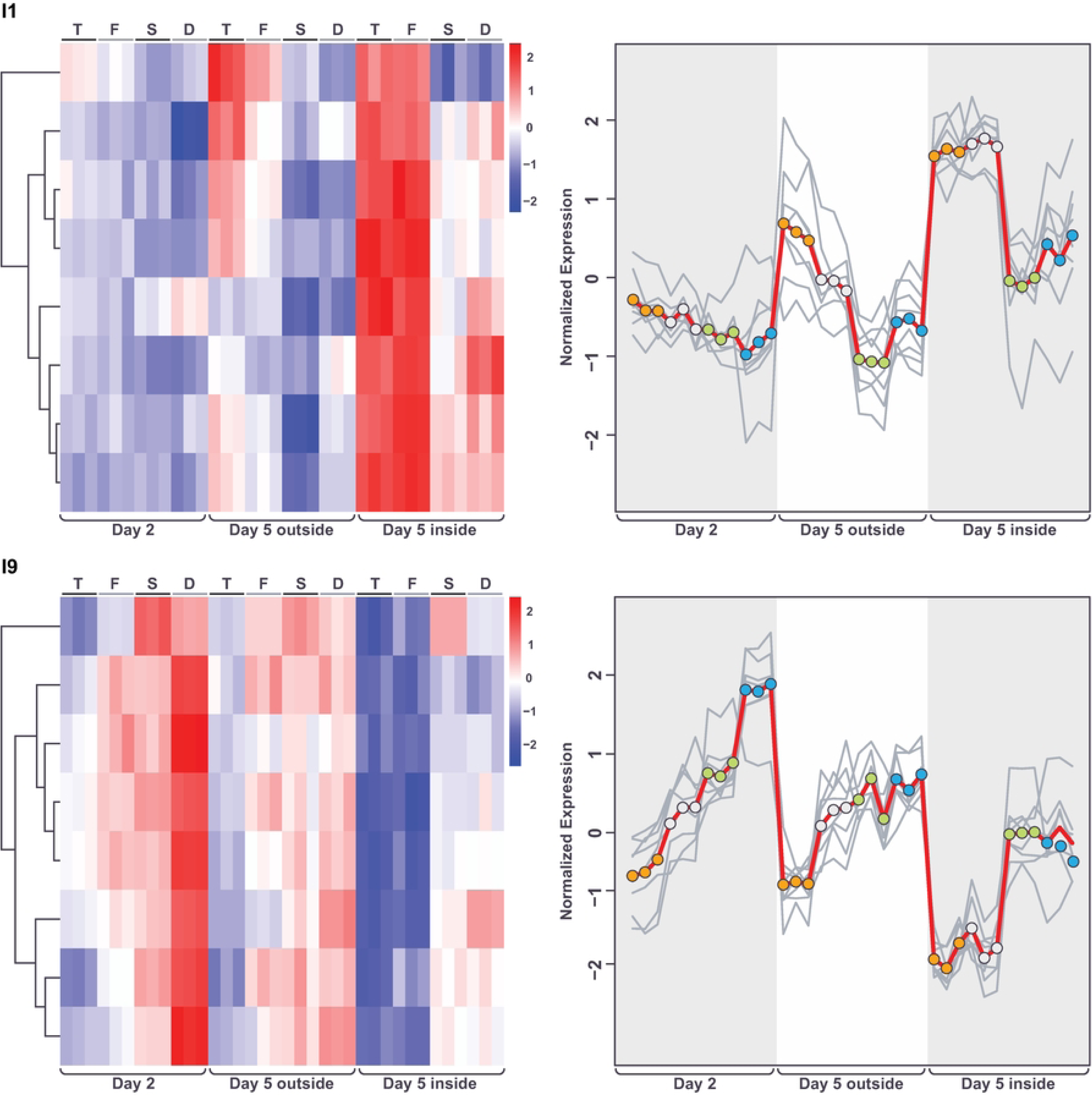
Heatmap showing hierarchical clustering of extracellular genes whose expression responds significantly to an additive model incorporating the genotype and spatiotemporal factors or whose expression indicates a significant non-additive interaction between the two factors. Color reflects log_2_ normalized expression for each gene (red high, blue low) and genotypes are indicated as T=*tec1*Δ, F=WT F13, S=*sfl1*Δ, D=*dig1*Δ.

**Figure 11.**
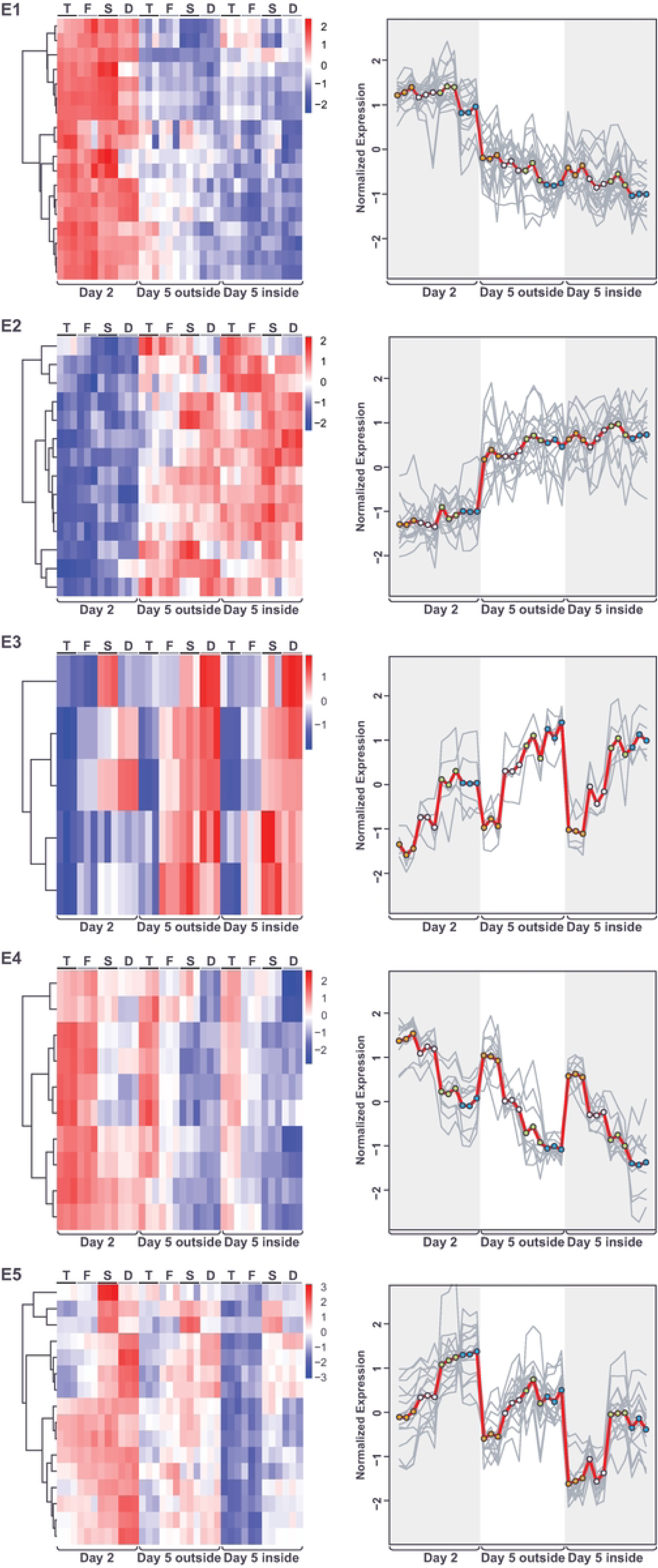
Gene expression patterns within the extracellular gene expression clusters E1-5 displayed as heatmaps (left) and expression traces (right) of all genes within each cluster (grey lines). In the expression traces, the mean expression of the genes in the cluster is highlighted in red line with circles reflecting genotype (orange=*tec1*Δ, grey=WT F13, green=*sfl1*Δ, blue=*dig1*Δ). Color in heatmaps reflects log_2_ normalized expression for each gene (red high, blue low) and genotypes are indicated as T=*tec1*Δ, F=WT F13, S=*sfl1*Δ, D=*dig1*Δ.

The genes responding mostly to the genotype factor fell into three main clusters. Two of these clusters showed patterns “reflected” for their behavior across the 4 genotypes in the day-5 samples, with expression higher in the *tec1*Δ and WT samples in cluster E4 and higher in the *sfl1*Δ and *dig1*Δ samples in cluster E5 (**Figure 11**). That is, the expression of these genes in the day-5 samples correlates positively or negatively with the degree of colony morphology seen in each genotype. Like clusters E1 and E2, genes encoding cell surface glycoproteins were found in both clusters E4 and E5, although these were more common in E4 (E4: *DAN4*, *HPF1*; E4: *CIS3*, *FLO10*, *FLO11*, *MSB2*, *PIR1*, *SRL1*) (**Table S17**). Cluster E4 also contained both of the genes encoding alpha-factor (*MF(ALPHA)1* and *MF(ALPHA)2*), *SAG1* and *AGA1* encoding alpha-agglutinin and the anchorage subunit of a-agglutinin, respectively, and several genes involved in cell wall remodelling during bud separation (*CTS1*, *DSE2*, DSE4). The third cluster mainly responding to genotype, E3, showed an expression pattern similar to E5 (**Figure 11**). This cluster consisted of a small number of cell wall genes (*CWP1*, *CWP2*, *SED1*, *PAU24*, *NFG1*), with 4/5 of these genes encoding known glycoproteins (the exception, *NFG1*, encodes a negative regulator of the filamentous growth pathway [34] (**Table S17**).

Looking more closely at clusters E4 and E5, it can be seen that the genes in each cluster vary in how closely their expression behavior at day-2 matches that at day-5. In each cluster, some genes show the same strong pattern of differential expression across genotypes at day-2 as they do at day-5, while other genes show a weaker version of this pattern and the remaining genes show little variation across genotypes at day-2 (**Figure 11**). These results are consistent with differential patterns of gene expression across genotypes that initiate at different timepoints in the development of the colony.

### Effect on Colony Morphology of Deleting Genes from Different Transcriptional Clusters

Our analysis so far identified a number of transcriptional clusters based on the effect of genotype and time/colony-region (spatiotemporal factor) on gene expression. Several of these clusters appear to have meaningful correlations with colony morphology, both positive and negative, for example clusters G1 and I9 demonstrate gene expression that correlates positively with the degree of colony structure associated with each genotype.

Based on this analysis we deleted 26 genes representing different clusters, with an emphasis on clusters that show correlations (positive or negative) with colony morphology. The genes that we chose to delete, and their cluster assignments are listed in **Table 3**.

**Table 3.** Genes deleted to test effect on colony morphology along with which cluster(s) they were assigned to.

| Systematic Name | Gene | Spatiotemporal<br>Cluster | Genotype<br>Cluster | Interaction<br>Cluster |
| --- | --- | --- | --- | --- |
| YPR194C | OPT2 | 0 | 0 | 1 |
| YHR143W | DSE2 | 0 | 0 | 3 |
| YJL078C | PRY3 | 0 | 0 | 3 |
| YGL028C | SCW11 | 0 | 0 | 3 |
| YBR093C | PHO5 | 0 | 0 | 4 |
| YLR042C | NFG1 | 0 | 0 | 4 |
| YIR019C | FLO11 | 0 | 0 | 9 |
| YGR014W | MSB2 | 0 | 0 | 9 |
| YKL043W | PHD1 | 0 | 0 | 9 |
| YIL123W | SIM1 | 0 | 0 | 9 |
| YOR247W | SRL1 | 0 | 0 | 9 |
| YPL163C | SVS1 | 0 | 0 | 9 |
| YKR013W | PRY2 | 0 | 1 | 0 |
| YLR286C | CTS1 | 0 | 2 | 0 |
| YJL160C | PIR5 | 0 | 2 | 0 |
| YGR041W | BUD9 | 1 | 2 | 0 |
| YOL155C | HPF1 | 1 | 2 | 0 |
| YKL096W | CWP1 | 2 | 1 | 0 |
| YLR353W | BUD8 | 3 | 1 | 0 |
| YOR315W | SFG1 | 3 | 1 | 0 |
| YJL158C | CIS3 | 5 | 1 | 0 |
| YIL162W | SUC2 | 5 | 1 | 0 |
| YNR067C | DSE4 | 5 | 2 | 0 |
| YNL327W | EGT2 | 5 | 2 | 0 |
| YNL066W | SUN4 | 5 | 2 | 0 |
| YGR066C | GID10 | 7 | 1 | 0 |

As expected from their known roles in development of complex colony morphologies [12, 25, 26], deletions of *FLO11* or MSB2, from cluster I9, produced colonies that remained fully smooth at day-5 (**Figure 12**). Deletion of other I9 genes tested (*PHD1*, *SIM1*, *SRL1* and *SVS1*) had little effect on F13 colony morphology, retaining a central smooth and outer structured zone at day-5. Among the remaining genes, deletion of *SFG1* (clusters C3+G1) and *CIS3* (clusters C5+G1) produced fully smooth colonies, while deletion of *BUD8* (clusters C3+G1) produced colonies with a smooth center and notably reduced structure in the outer region of the colony. The remaining deletions had little effect on F13 colony morphology (**Figure 12**).

**Figure 12.**
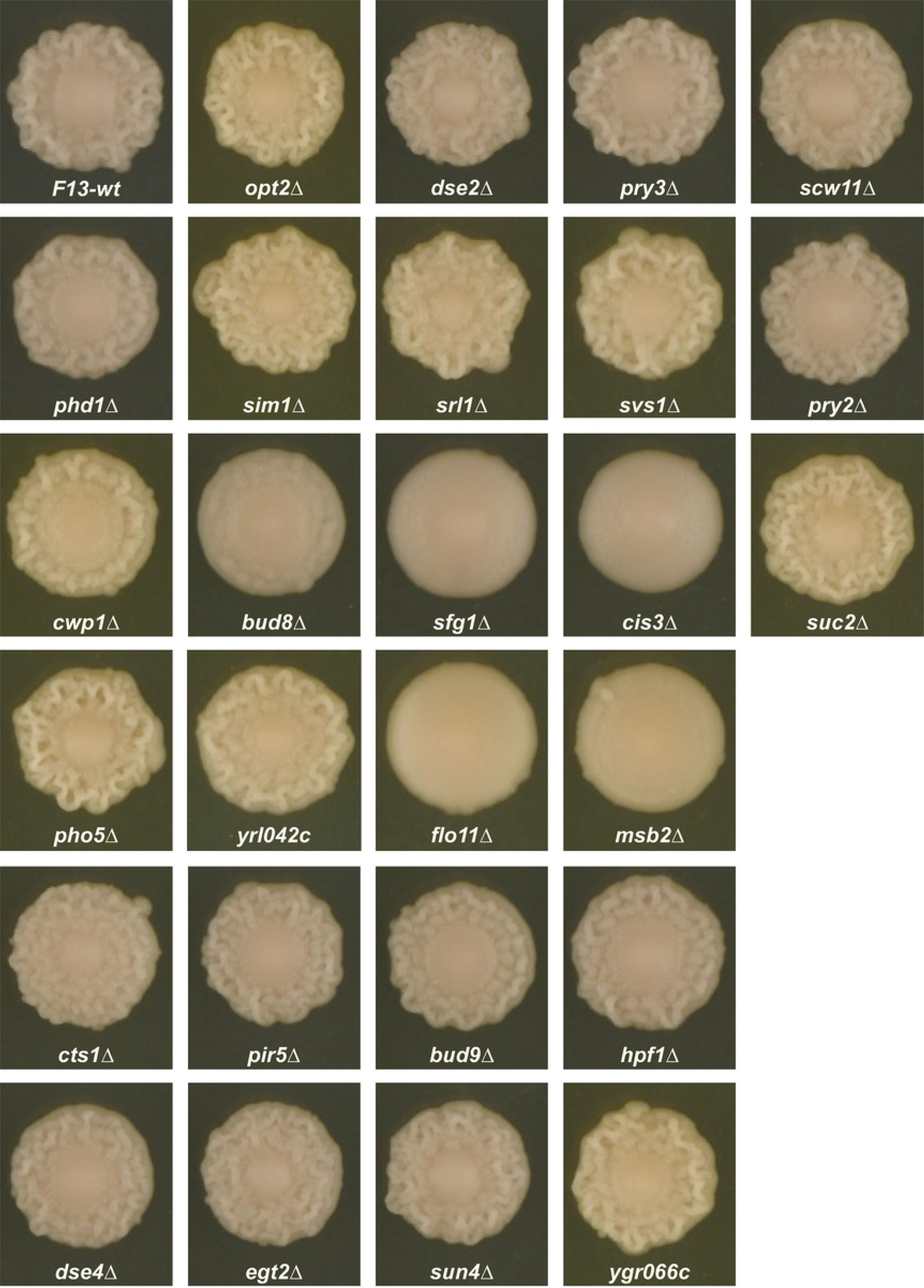
Effect on day-5 F13 colony morphology of deleting genes from a range of clusters (listed in **Table 3**).

## Discussion

Our study identified a number of coherent gene clusters in which expression varied by colony age and region and in the presence of genetic perturbations increasing or decreasing the degree of colony morphology. The major expression patterns associated with the genotype factor showed a striking positive or negative correlation with the degree of colony morphology achieved by colonies of each genotype on day-5. The major expression patterns associated with the spatiotemporal factor showed a strong difference between the day-2 and day-5-inside samples with the day-5-outside sample either resembling one of the other two, or displaying an intermediate expression level. Interestingly, some strong overlaps were seen between the genotype and spatiotemporal clusters, with an over-representation of genes that are more highly expressed in the structured genotypes and are also more highly expressed at day-5, when colonies can become fully ruffled, than at day-2, before colony structure has fully developed. A similar overlap was seen between the complementary clusters with highest expression in the unstructured genotypes and having highest expression at day-2.

Surprisingly, the genes positively correlated with the ruffled phenotype (by genotype and colony age/region) and those negatively correlated were enriched for many of the same GO terms, with over-representation of cell wall / extracellular genes in both directions. This suggests that extensive cell wall composition changes accompany changes in colony morphology, with numerous extracellular genes up- and down-regulated. When we looked specifically at the expression pattern of extracellular genes this pattern was confirmed, with some genes most highly expressed in the most structured genotypes and others most highly expressed in the smooth genotypes.

Although the genes in each of these two clusters showed consistent expression behavior in the day-5 samples, the behavior of the genes at day-2 varied within each cluster. For some genes, the pattern of differential expression across genotypes that was seen in the day-5 samples was also seen at day-2, whereas for other genes this pattern was weaker or absent. This suggests that the time at which the pattern of differential expression first develops varies between genes. By day-5 all of these genes show differential expression across genotypes, largely correlating with the degree of colony structure observed at that time, but some genes are already strongly differentially expressed by day-2, when the *tec1*Δ and WT colonies are fully smooth and the *sfl1*Δ and *dig1*Δ colonies have only begun to show weak signs of colony structure. The genes showing early differential expression may be important for specifying the development of colony structure at later time points, i.e. they may be part of a “developmental program” for colony structure. In contrast, genes showing differential expression only after the full development of colony morphology may be physical effectors of the phenotype, or genes responding to different “environmental” conditions within ruffled versus smooth colonies.

Deletion of a set of genes sampled from different clusters only identified 5 with strong effects on colony morphology (*BUD8*, *CIS3*, *FLO11*, *MSB2* and *SFG1*), all of which greatly reduced or eliminated the structure of the F13 outer region. These 5 genes are quite diverse: *FLO11* and *MSB2* encode cell surface glycoproteins with known roles in promoting colony structure [12, 25, 26]; similarly, *CIS3* encodes a cell wall glycoprotein whose deletion causes loss of colony morphology in strain F45, related to the F13 strain used here [16]; *SFG1* encodes a transcription factor known to promote superficial pseudohyphae and invasive growth [34, 35], *BUD8* is involved in bud site selection [36]. While our analysis was by no means comprehensive, it is interesting that we did not observe any genes whose deletion caused F13 to become fully ruffled. This is consistent with the outer, structured region of F13 representing a more specialized and elaborate developmental pathway than the smooth interior. Deletion of any of the genes promoting or effecting the outer colony morphology would then produce fully smooth colonies. In contrast, under this model only deletions of negative regulators of the ruffled developmental program, such as *SFL1* or *DIG1*, would produce fully structured colonies.

The expression changes observed in our study were dominated by those associated with the spatiotemporal factor. The major clusters associated with this factor consistently showed a strong difference between the day-2 and day-5-inside samples, but were distinguished by the behavior of the day-5-outside sample. In some clusters the day-5 outside sample behaved like either the day-2 or day-5-inside samples, while in others it showed an expression pattern intermediate between the other two, i.e. genes showing unique expression in the day-5 outside samples were rare. Instead, the patterns that we do observe are consistent with gene expression in response to a buildup of signaling molecules or metabolic waste products within the colony over time, or conversely depletion of nutrients within the colony and adjacent regions of agar. Gene expression responding to these signals would place the day-2 sample and the day-5-inside sample at opposite extremes, with the day-5-outside sample showing intermediate expression, with its position relative to the other two depending on the steepness of concentration gradient(s) associated with the chemical(s) controlling gene expression, and any thresholds that exist for response to those gradients. Similarly, the oldest cells in the day-5 colony are likely to be in the inside region and the youngest in the outside region [37], so that average cell age could also vary from day-2 (youngest) to the day-5-inside sample (oldest) with the day-5-outside sample intermediate between the other two.

Several previous studies have identified patterns of spatial cell differentiation within yeast colonies. As well as differentiation within structured colonies [6, 7], work from Palkova and colleagues has also identified patterns of vertical and horizontal cell differentiation within old smooth colonies. Such colonies display stratification into clearly demarcated upper (U) and lower (L) cell populations, with distinctive gene expression patterns [38], while programmed cell death is confined to the central region of old colonies [37]. The sharp demarcation between the U and L populations is similar to the clearly spatial defined subpopulation of sporulating cells observed by Honigberg and colleagues within diploid yeast colonies [39].

One interesting question raised by our work and previous studies is the precise nature of the signals that specify spatial (and temporal) differentiation within yeast colonies. The sporulating region within diploid colonies has been shown to reflect an overlap between subpopulations expressing the *IME1* and *IME2* activators of sporulation and to respond to alkaline pH signaling through the Rim101/PacC pathway [39]. Similarly, the localization of programmed cell death to the inner regions of old colonies has been shown to be dependent on ammonia signaling [37]. However, the precise nature of the full set of environmental and cell-cell signals specifying patterns of yeast colony differentiation have yet to be fully elucidated in any system.

Our work described here extends previous studies on yeast colony development to examine the interaction between genetic and spatiotemporal factors in the development of a highly structured colony morphology. Our results suggest a complex interplay between genetic perturbations and their effect on spatially defined subpopulations within a biofilm-model.

## Supplementary Material

File S1. S288c reference FASTA sequence, extended to include genes present in F45 and not S288c.

File S2. S288c reference GFF file, extended to include non-coding RNAs (ncRNAs) and genes present in F45 and not S288c.

File S3. Raw read counts and normalized log_2_ read counts per gene per library.

File S4. R script used for data processing and analysis.

Table S1. Functional enrichment analysis for genes in cluster G1.

Table S2. Functional enrichment analysis for genes in cluster G2.

Table S3. Functional enrichment analysis for genes in cluster G3.

Table S4. Functional enrichment analysis for genes in cluster C1.

Table S5. Functional enrichment analysis for genes in cluster C2.

Table S6. Functional enrichment analysis for genes in cluster C3.

Table S7. Functional enrichment analysis for genes in cluster C4.

Table S8. Functional enrichment analysis for genes in cluster C5.

Table S9. Functional enrichment analysis for genes in cluster C6.

Table S10. Functional enrichment analysis for genes in cluster C7.

Table S11. Functional enrichment analysis for genes in cluster C8.

Table S12. Functional enrichment analysis for genes found in both clusters G1 and C2.

Table S13. Functional enrichment analysis for genes found in both clusters G2 and C1.

Table S14. Functional enrichment analysis for all genes showing significant non-additive interaction between the genotype and spatiotemporal factors.

Table S15. Functional enrichment analysis for genes in cluster I9.

Table S16. Functional enrichment analysis for genes in cluster I1.

Table S17. Genes in extracellular clusters E1-5.

## Notes

### Competing Interest Statement

The authors have declared no competing interest.

